# *pipesnake*: Generalized software for the assembly and analysis of phylogenomic datasets from conserved genomic loci

**DOI:** 10.1101/2024.02.13.580223

**Authors:** Ian G. Brennan, Sonal Singhal, Ziad Al Bkhetan

## Abstract

**Motivation:** Phylogenetics has moved into the era of genomics, incorporating enormous volumes of data to study questions at both shallow and deep scales. With this increase in information, phylogeneticists need new tools and skills to manipulate and analyze these data. To facilitate these tasks and encourage reproducibility, the community is increasingly moving towards automated workflows.

**Results:** Here we present pipesnake, a phylogenomics pipeline written in Nextflow for the processing, assembly, and phylogenetic estimation of genomic data from short-read sequences. pipesnake is an easy to use and efficient software package designed for this next era in phylogenetics.

**Availability:** The quick brown fox jumps over the lazy dog.

**Supplementary information:** pipesnake is written in Nextflow and freely available from https://github.com/AusARG/pipesnake alongside comprehensive documentation and a wiki/tutorial. The pipeline is containerized and deployable via Docker, Singularity, and Conda making it easy to run on different compute infrastructures.

## 1 Introduction

Genomic datasets employing hundreds to tens of thousands of individual markers have become common across phylogenetics, helping to resolve questions from intra-specific to inter-class scales. These projects typically target genomic loci which can be reliably identified, aligned, and modeled, and are sufficiently conserved that they remain useful at varied phylogenetic depths. Popular examples of these marker sets include Anchored Hybrid Enrichment (AHE—Lemmon et al. 2012) and Ultra Conserved Elements (UCE—Faircloth et al. 2012), however, new marker sets are being designed regularly for use at both broad and narrow foci (Johnson et al. 2019—Angiosperms353; Hutter et al. 2022—FrogCap; Hughes et al. 2018—FishLife). The primary impediment to developing these resources is the necessary bioinformatics architecture to execute downstream steps after data generation, including: raw read filtering and trimming, sequence read assembly, contig mapping, orthology assignment, target sequence collation, alignment and alignment processing (quality assessment, filtering, trimming), locus-specific phylogenetic estimation, and species tree estimation. With the notable exception of PHYLUCE (Faircloth 2016)— which provided the inspiration for this work—most solutions are bespoke, proprietary, or poorly documented. Many limitations to the broader use of such analysis workflows are due to clade or marker set specifics, or software installations that make reuse onerous. Here we introduce the software package *pipesnake* which has been designed to handle varied data and target types, and executed in Python and Nextflow computer languages for simplicity of data handling.

### 2 The *pipesnake* Workflow

*pipesnake* is a workflow intended to take a batch of data from second-generation short-read sequences to a species tree estimate while retaining valuable intermediate files. In the most basic form, the user provides a comma-separated sample input file, a fasta file of target loci, and raw sequence read files. From the sample info file *pipesnake* identifies all reads corresponding to a specific sample (forward and reverse), identifies and concatenates reads from multiple lanes (if applicable), and passes them to BBMAP (Bushnell 2014) to remove duplicate reads (dedupe.sh). Deduplicated reads are then submitted to TRIMMOMATIC (Bolger et al. 2014) for residual adapter and barcode removal and set into read pairs using PEAR (Zhang et al. 2014). Trimmed and paired reads are then optionally passed to BBMAP for mapping against phylogenetically informed target sequences to remove off-target reads prior to assembly (which may otherwise slow the assembly process). pipesnake relies on TRINITY (Grabherr et al. 2011) or SPAdes (Prjibelski et al. 2020) for contig assembly, after which contigs are mapped to target sequences via BLAT (Kent 2002) via a reciprocal search. Highest quality contig-to-target matches are then extracted and pulled into a sample-specific fasta file (which we call a Pseudo-Reference Genome—PRG). The program then pulls each target locus into a marker-specific raw alignment before passing these files along to MAFFT (Katoh & Standley 2013) for initial alignment. Alignments can then be optionally refined and trimmed using GBLOCKS (Talavera & Castresana 2007). Final alignments are then used as input for phylogenetic estimation using the preferred software (RAxML—Stamatakis 2014; or IQTREE—Minh et al. 2020). Locus trees are collated and used as input for ASTRAL (Zhang et al. 2022) to estimate a species tree. A simplified diagram of the pipeline is included in Figure 1.

**Figure 1.**
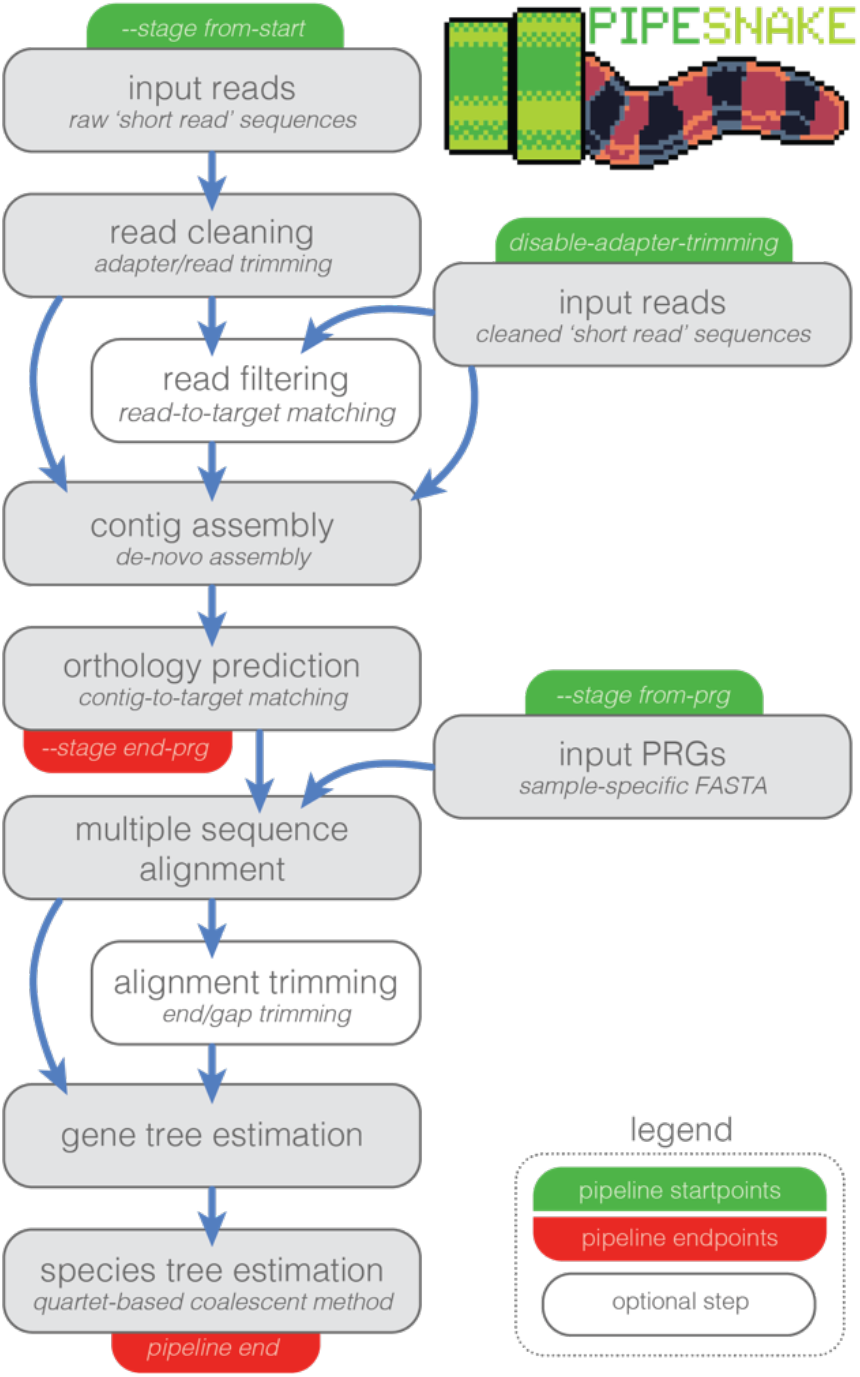
Simplified schematic of pipesnake workflow highlighting the various starting and stopping points and order of pipeline processes including optional steps.

At each step the *pipesnake* workflow generates a number of output files that are stored in process-specific directories. This allows the user to store and inspect intermediate files such as individual sample PRGs, alignment files, and locus trees. The modularity of the workflow means that if interrupted, rerunning *pipesnake* simply picks up where it left off (-resume), without the need to begin from scratch, and without the need to specify which step to begin at. Modularity also means that modifying the workflow is relatively straightforward. For example if a user would prefer to assemble contigs with software other than SPAdes or TRINITY, this requires only creating a new module file to pull a container of the software and specifying the process, outlining default parameters under docs/usage.md file, and fitting the new module into the workflow chronology in the primary pipesnake.nf file.

### 3 Motivation and Example

Squamate reptiles are the largest living non-fish vertebrate clade (11500+ species), spread widely across the globe from deserts to jungles, mountain peaks to oceanic islands, and nearly everywhere in-between. In addition to their incredible species richness and morphological diversity, squamate systematics have been an early focus for phylogenomics (Townsend et al. 2008; Wiens et al. 2010; Mulcahy et al. 2012). Dozens of empirical studies have used UCEs and AHE to investigate the squamate tree of life at both shallow and deep scales (Streicher & Wiens 2017; Brandley et al. 2015; Burbrink et al. 2019). To unify these marker sets Singhal et al. (2017) developed the Squamate Conserved Loci kit (SqCL) (Singhal et al. 2017) which incorporates more than 5400 genomic targets (∼5k UCEs, ∼400 AHE, ∼40 ‘legacy’ exons). Until now, the SqCL marker set has not had a well-documented reproducible workflow for assembling and analyzing new data. The lack of a user-friendly workflow has potentially acted as a limitation towards adopting this marker set, and so we present a reliable and consistent approach here. *pipesnake* is not, however, limited to assembling and analyzing SqCL data. The pipeline can also be comfortably used to assemble target loci from other organismal groups and for other shortread sequence data such as transcriptomes and genomes.

### 4 Implementation, Flexibility, Reproducibility

*pipesnake* is written in Nextflow which allows a flexible and easily customizable workflow execution on various compute infrastructures via Docker, Singularity or Conda packages. Our motivation for building *pipesnake* in Nextflow was so that we could easily provide support for local and HPC based executions. This allows *pipesnake* to interact seamlessly with workload managers like SLURM to optimize performance and juggle individual jobs. On initial use the workflow pulls necessary dependencies from online resources such as the Galaxy repository, Quay, and Bioconda. On future uses the workflow checks for locally cached software containers, excluding the need for manual local installation of dependencies. To take advantage of available resources, memory and CPU usage can be adapted by the user (see conf/base.config) and passed to pipesnake (e.g. -profile or -c) to optimize efficiency.

Regarding flexibility, many elements of the workflow’s behavior can be fine-tuned using a common syntax (--[process]_args) followed by processspecific arguments between two quotes. For example, specifying a given substitution model for IQTREE (e.g. --iqtree_args “-m GTR”) is trivial and additional process-specific arguments can be strung together. Parameter defaults are stored in the conf/base.config file and specifics about their application and usage can be found in docs/usage.md. To facilitate the synthesis of separate phylogenomics projects the *pipesnake* workflow can be initiated from the alignment-formation step using the --stage command (e.g. --stage from-prg). In this instance the user provides a comma-separated sample input file and paths to PRGs of interest in fasta format, avoiding the need to reassemble samples from raw data and eliminating computational burden.

To encourage transparent and reproducible methods, *pipesnake* generates a pipeline information file that stores the versions of all software used, in addition to reports on memory and CPU usage, Nextflow commands executed, specified parameters, and a complete log file. This design means the full pipeline can be run from a single command and rerun under the same or new parameters easily. We include an example dataset of four samples which under default parameters runs from raw sequence reads to an output species tree in just a matter of minutes on a local desktop machine. Instructions for using *pipesnake*—from installation to running it on your own data—are available in the wiki/tutorial at https://github.com/AusARG/pipesnake/wiki.

### 5. Performance Evaluation

To provide evidence of performance, we test the pipeline on a dataset of 9 samples with gzip compressed paired-end fastq files with an average size of 208 MB (min 1.3MB and max is 624M). Read files comprise 150 bp paired-end reads generated from SqCL hybrid enrichment across 5400 targets of varied length. Details on the required resources to run this workflow are reported in Fig.S1 for memory and time, respectively. In this example, the assembler TRINITY is the bottleneck when it comes to the performance time required to finish the analysis. TRINITY and BBMAP deduplication require more memory than other processes. However, the required resources are within the computing power available to many genomics researchers.

## Acknowledgements

The authors acknowledge the provision of computing and data resources provided by the Australian BioCommons Leadership Share (ABLeS) program and Nextflow Tower service. These programs are co-funded by Bioplatforms Australia (enabled by NCRIS), the National Computational Infrastructure and Pawsey Supercomputing Centre. Special thanks to Sophie Mazard and Johan Gustafsson for initiating this fruitful collaboration, and to Scott Keogh, Damien Esquérre, and Sarin Tiatragul for product testing and support.

## Supplementary Figures

**Figure S1.**
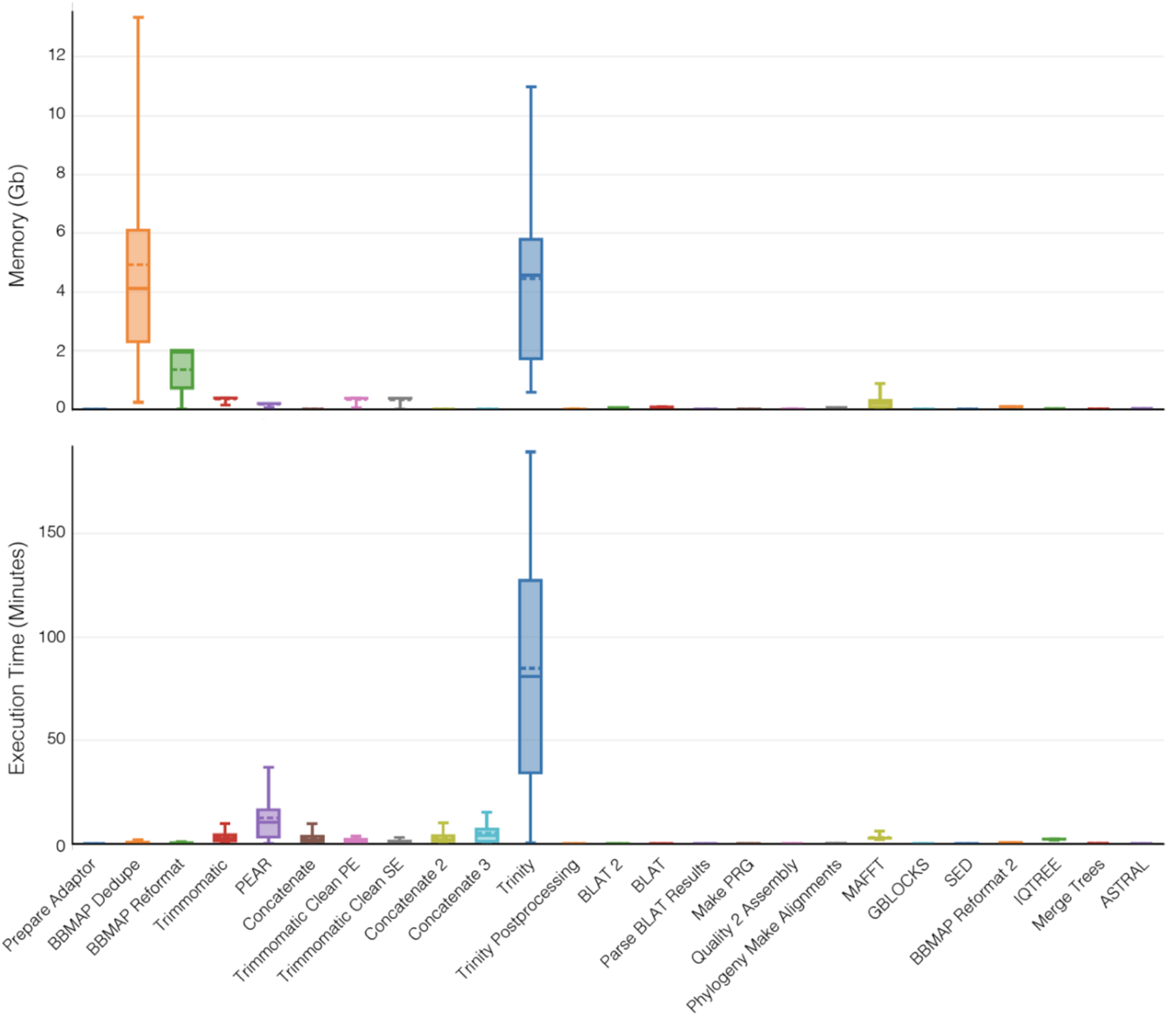
Memory requirements (top) and execution time (bottom) for each step in the workflow as carried out on nine example samples. Steps are ordered chronologically. Box plots reflect all nine samples up until the Phylogeny Make Alignments step, at which point they were batched into nine batches of 250 alignment files and passed to the remaining steps. As shown, TRINITY is the primary bottleneck for performance time.

## References

Bankevich, A., Nurk, S., Antipov, D., Gurevich, A. A., Dvorkin, M., Kulikov, A. S., & Pevzner, P. A. (2012). SPAdes: a new genome assembly algorithm and its applications to single-cell sequencing. Journal of computational biology, 19(5), 455–477.

Bolger, A. M., Lohse, M., & Usadel, B. (2014). Trimmomatic: a flexible trimmer for Illumina sequence data. Bioinformatics, 30(15), 2114–2120.

Brandley, M. C., Bragg, J. G., Singhal, S., Chapple, D. G., Jennings, C. K., Lemmon, A. R., … & Moritz, C. (2015). Evaluating the performance of anchored hybrid enrichment at the tips of the tree of life: a phylogenetic analysis of Australian Eugongylus group scincid lizards. BMC Evolutionary Biology, 15(1), 1–14.

Burbrink, F. T., Grazziotin, F. G., Pyron, R. A., Cundall, D., Donnellan, S., Irish, F., … & Zaher, H. (2020). Interrogating genomic-scale data for Squamata (lizards, snakes, and amphisbaenians) shows no support for key traditional morphological relationships. Systematic biology, 69(3), 502–520.

Bushnell, B. (2014). BBMap: a fast, accurate, splice-aware aligner (No. LBNL-7065E). Lawrence Berkeley National Lab.(LBNL), Berkeley, CA (United States).

Faircloth, B. C. (2016). PHYLUCE is a software package for the analysis of conserved genomic loci. Bioinformatics, 32(5), 786–788.

Faircloth, B. C., McCormack, J. E., Crawford, N. G., Harvey, M. G., Brumfield, R. T., & Glenn, T. C. (2012). Ultraconserved elements anchor thousands of genetic markers spanning multiple evolutionary timescales. Systematic biology, 61(5), 717–726.

Grabherr, M. G., Haas, B. J., Yassour, M., Levin, J. Z., Thompson, D. A., Amit, I., … & Regev, A. (2011). Trinity: reconstructing a full-length transcriptome without a genome from RNA-Seq data. Nature biotechnology, 29(7), 644.

Hughes, L. C., Ortí, G., Huang, Y., Sun, Y., Baldwin, C. C., Thompson, A. W., … & Shi, Q. (2018). Comprehensive phylogeny of ray-finned fishes (Actinopterygii) based on transcriptomic and genomic data. Proceedings of the National Academy of Sciences, 115(24), 6249–6254.

Hutter, C. R., Cobb, K. A., Portik, D. M., Travers, S. L., Wood Jr, P. L., & Brown, R. M. (2022). FrogCap: A modular sequence capture probe-set for phylogenomics and population genetics for all frogs, assessed across multiple phylogenetic scales. Molecular Ecology Resources, 22(3), 1100–1119.

Johnson, M. G., Pokorny, L., Dodsworth, S., Botigué, L. R., Cowan, R. S., Devault, A., … & Wickett, N. J. (2019). A universal probe set for targeted sequencing of 353 nuclear genes from any flowering plant designed using k-medoids clustering. Systematic biology, 68(4), 594–606.

Katoh, K., & Standley, D. M. (2013). MAFFT multiple sequence alignment software version 7: improvements in performance and usability. Molecular biology and evolution, 30(4), 772–780.

Kent, W. J. (2002). BLAT—the BLAST-like alignment tool. Genome research, 12(4), 656–664.

Lemmon, A. R., Emme, S. A., & Lemmon, E. M. (2012). Anchored hybrid enrichment for massively high-throughput phylogenomics. Systematic Biology, 61(5), 727–744.

Minh, B. Q., Schmidt, H. A., Chernomor, O., Schrempf, D., Woodhams, M. D., Von Haeseler, A., & Lanfear, R. (2020). IQ-TREE 2: new models and efficient methods for phylogenetic inference in the genomic era. Molecular biology and evolution, 37(5), 1530–1534.

Mulcahy, D. G., Noonan, B. P., Moss, T., Townsend, T. M., Reeder, T. W., Sites Jr, J. W., & Wiens, J. J. (2012). Estimating divergence dates and evaluating dating methods using phylogenomic and mitochondrial data in squamate reptiles. Molecular Phylogenetics and Evolution, 65(3), 974–991.

Prjibelski, A., Antipov, D., Meleshko, D., Lapidus, A., & Korobeynikov, A. (2020). Using SPAdes de novo assembler. Current protocols in bioinformatics, 70(1), e102.

Singhal, S., Grundler, M., Colli, G., & Rabosky, D. L. (2017). Squamate conserved loci (Sq CL): a unified set of conserved loci for phylogenomics and population genetics of squamate reptiles. Molecular ecology resources, 17(6), e12–e24.

Stamatakis, A. (2014). RAxML version 8: a tool for phylogenetic analysis and post-analysis of large phylogenies. Bioinformatics, 30(9), 1312–1313.

Streicher, J. W., & Wiens, J. J. (2017). Phylogenomic analyses of more than 4000 nuclear loci resolve the origin of snakes among lizard families. Biology Letters, 13(9), 20170393.

Talavera, G., & Castresana, J. (2007). Improvement of phylogenies after removing divergent and ambiguously aligned blocks from protein sequence alignments. Systematic biology, 56(4), 564–577.

Townsend, T. M., Alegre, R. E., Kelley, S. T., Wiens, J. J., & Reeder, T. W. (2008). Rapid development of multiple nuclear loci for phylogenetic analysis using genomic resources: an example from squamate reptiles. Molecular phylogenetics and evolution, 47(1), 129–142.

Wiens, J. J., Kuczynski, C. A., Townsend, T., Reeder, T. W., Mulcahy, D. G., & Sites Jr, J. W. (2010). Combining phylogenomics and fossils in higher-level squamate reptile phylogeny: molecular data change the placement of fossil taxa. Systematic Biology, 59(6), 674–688.

Zhang, J., Kobert, K., Flouri, T., & Stamatakis, A. (2014). PEAR: a fast and accurate Illumina Paired-End reAd mergeR. Bioinformatics, 30(5), 614–620.

Zhang, C., & Mirarab, S. (2022). Weighting by gene tree uncertainty improves accuracy of quartet-based species trees. Molecular Biology and Evolution, 39(12), msac215.

